## Supplementary Materials for "Evaluation of SARS-CoV-2 Entry, Inflammation and New Therapeutics in Human Lung Tissue Cells"

### Figure S1

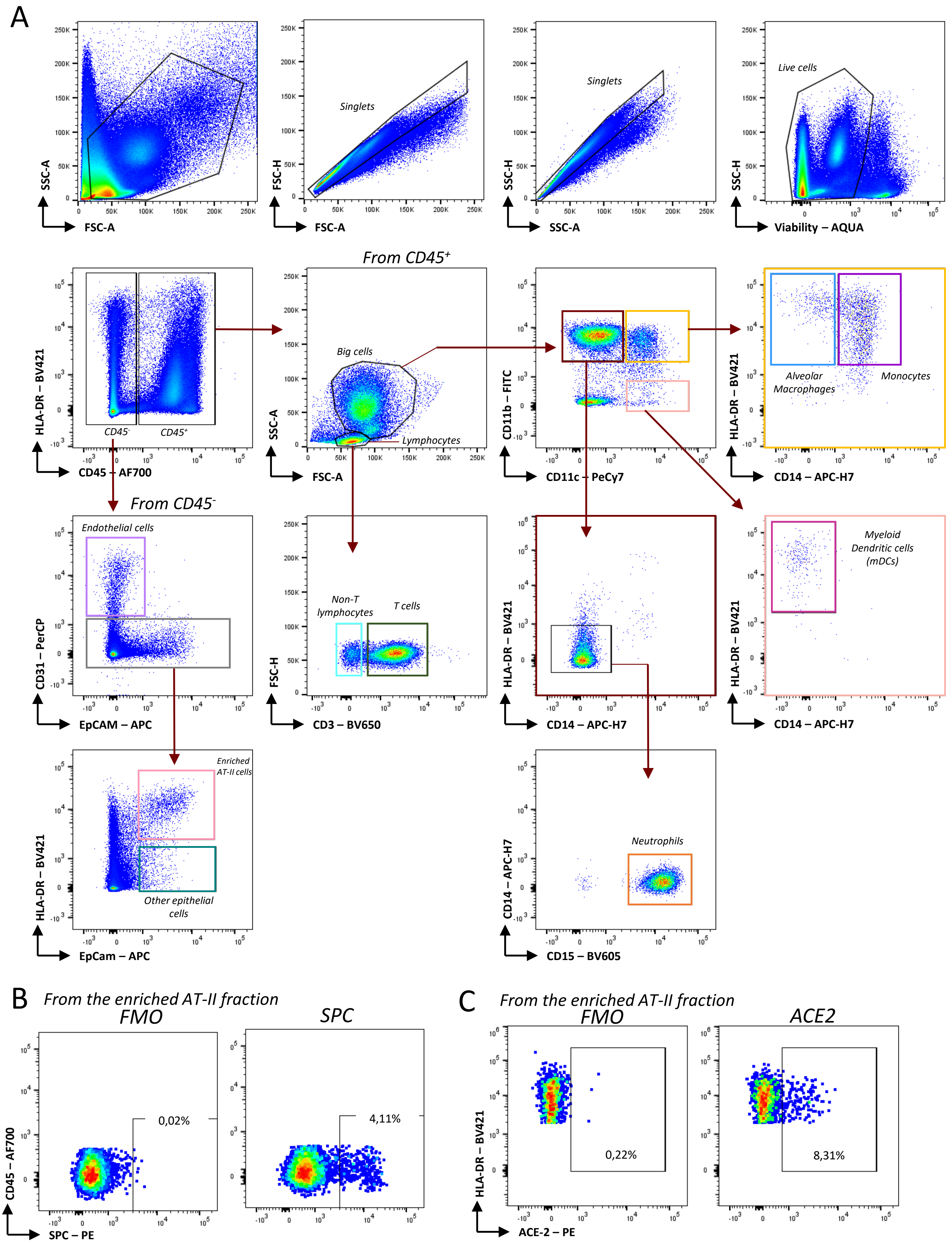
A
SSC-A
FSC-A
FSC-H
Singlets
SSC-H
Live cells
Viability – AQUA
From CD45+
HLA-DR – BV421
CD45-
CD45+
CD45 – AF700
Big cells
Lymphocytes
CD11b – FITC
CD11c – PeCy7
Alveolar Macrophages
Monocytes
CD14 – APC-H7
From CD45-
Endothelial cells
CD31 – PerCP
EpCAM – APC
Non-T lymphocytes
T cells
CD3 – BV650
Myeloid Dendritic cells (mDCs)
Enriched AT-II cells
Other epithelial cells
EpCam – APC
Neutrophils
CD14 – APC-H7
CD15 – BV605
B
From the enriched AT-II fraction
FMO
SPC
0,02%
4,11%
CD45 – AF700
SPC – PE
C
From the enriched AT-II fraction
FMO
ACE2
0,22%
8,31%
HLA-DR – BV421
ACE-2 – PE

### Figure S2

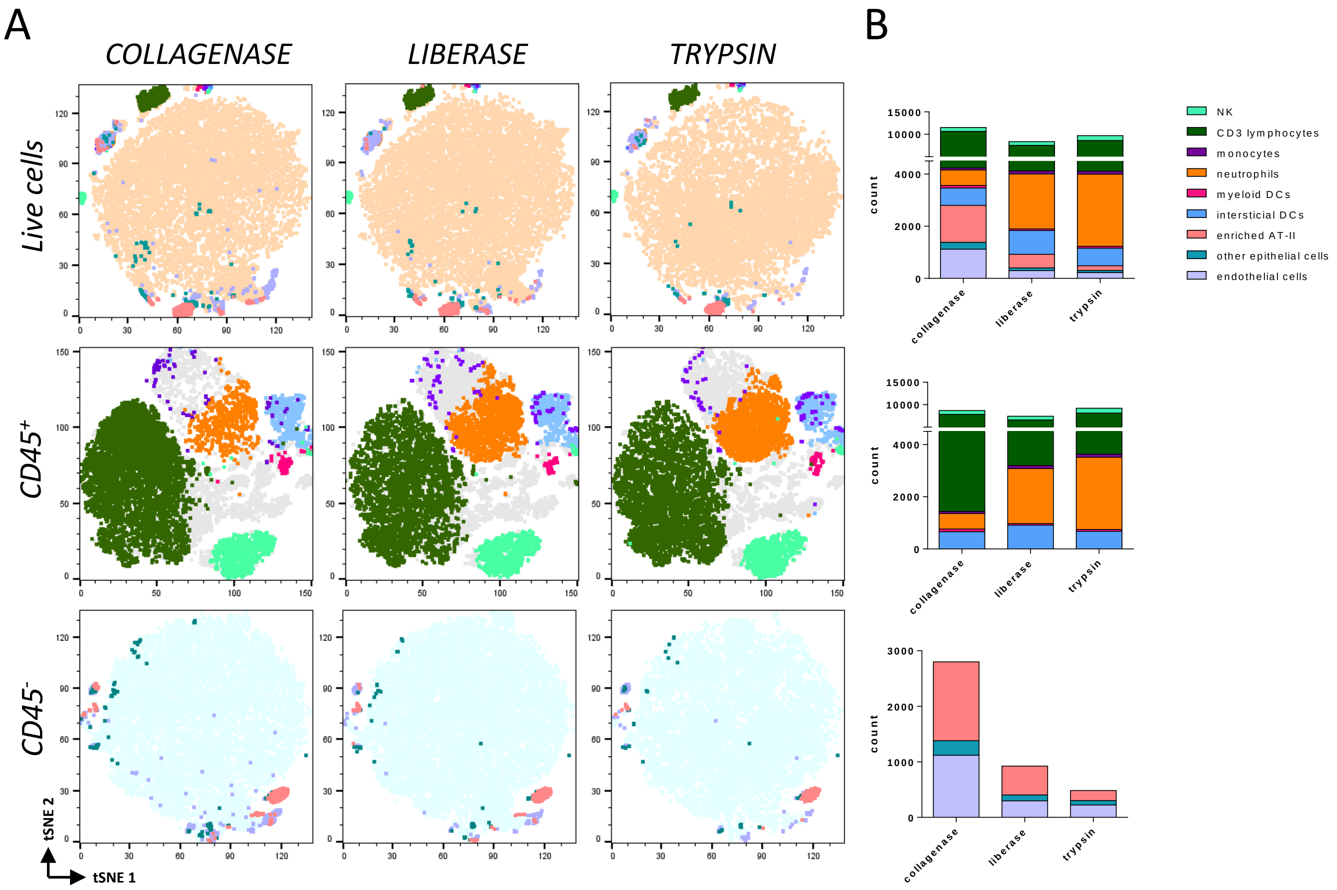
A
B
COLLAGENASE
LIBERASE
TRYPSIN
Live cells
CD45+
CD45-
tSNE 2
tSNE 1
NK
CD3 lymphocytes
monocytes
neutrophils
myeloid DCs
intersticial DCs
enriched AT-II
other epithelial cells
endothelial cells
count
collagenase
liberase
trypsin

### Figure S3

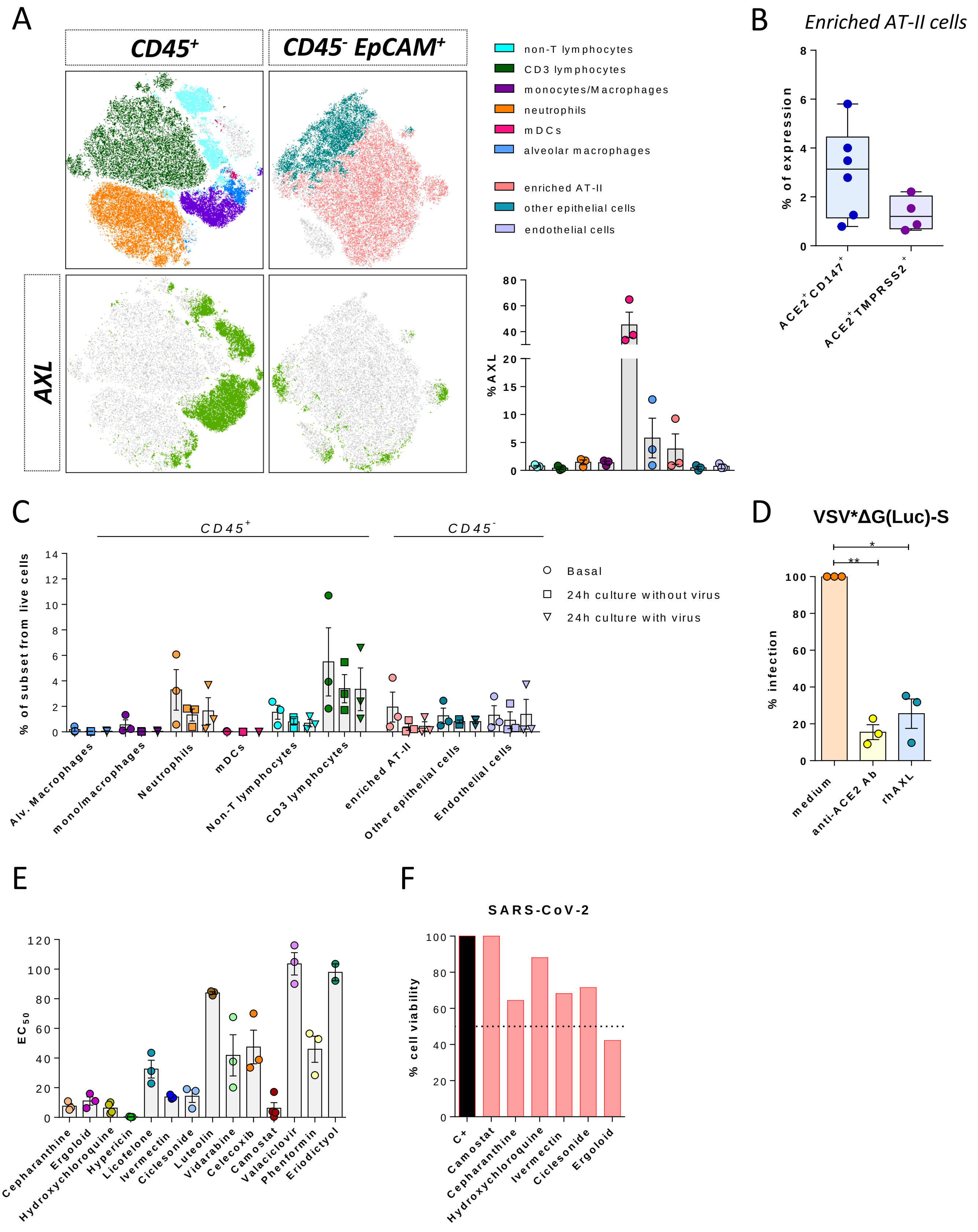
A
CD45+
CD45- EpCAM+
AXL
non-T lymphocytes
CD3 lymphocytes
monocytes/Macrophages
neutrophils
mDCs
alveolar macrophages
enriched AT-II
other epithelial cells
endothelial cells
% AXL
B
Enriched AT-II cells
% of expression
ACE2+CD147+
ACE2+TMPRSS2+
C
CD45+
CD45-
% of subset from live cells
Basal
24h culture without virus
24h culture with virus
Alv. Macrophages
mono/macrophages
Neutrophils
mDCs
Non-T lymphocytes
CD3 lymphocytes
enriched AT-II
Other epithelial cells
Endothelial cells
D
VSV*ΔG(Luc)-S
% infection
medium
anti-ACE2 Ab
rhAXL
E
EC50
Cepharanthine
Ergoloid
Hydroxychloroquine
Hypericin
Licofelone
Ivermectin
Ciclesonide
Luteolin
Vidarabine
Celecoxib
Camostat
Valaciclovir
Phenformin
Eriodictyol
F
SARS-CoV-2
% cell viability
C+
Camostat
Cepharanthine
Hydroxychloroquine
Ivermectin
Ciclesonide
Ergoloid

### Figure S4

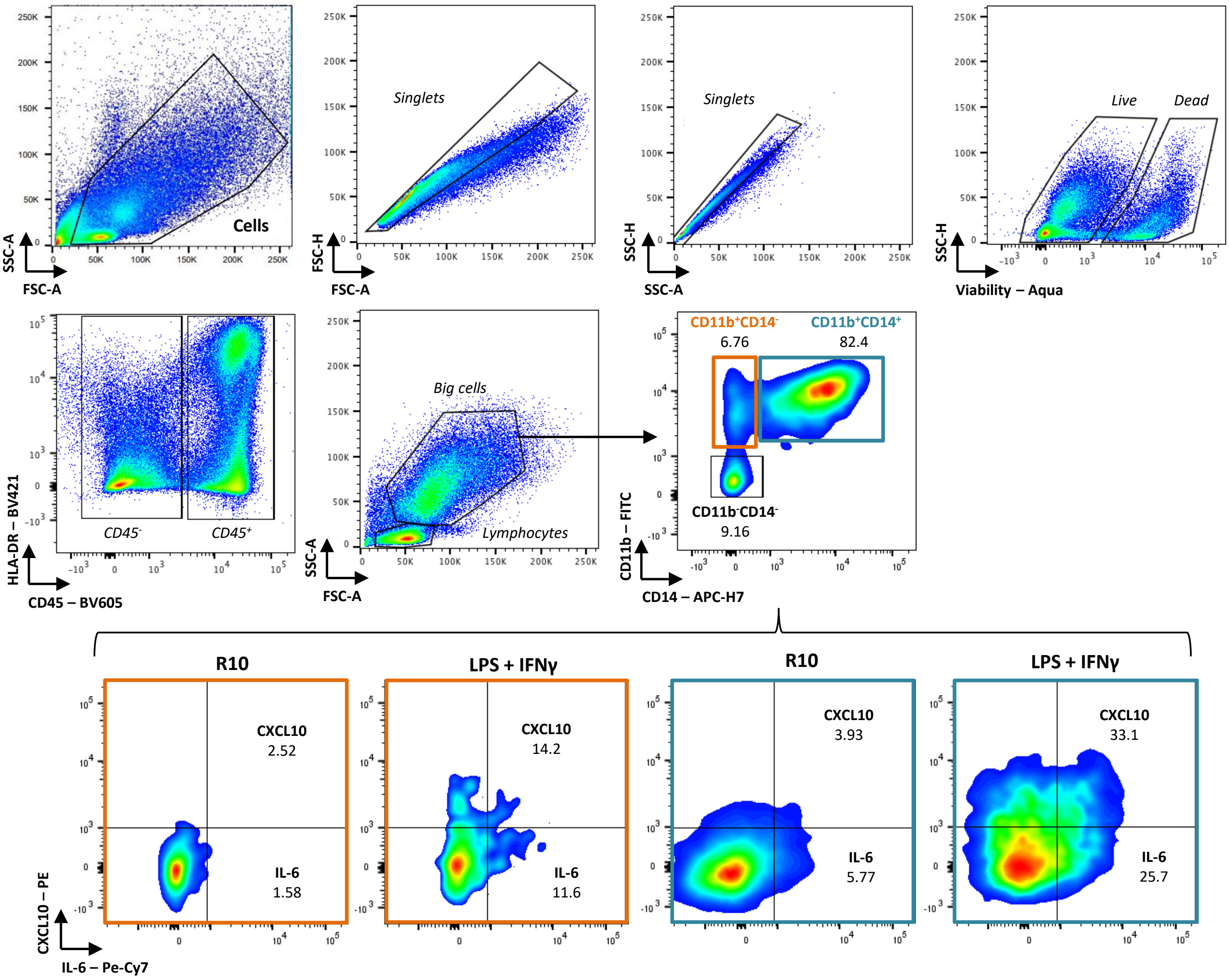

Cells
SSC-A
FSC-A
Singlets
FSC-H
FSC-A
Singlets
SSC-H
SSC-A
Live
Dead
SSC-H
Viability – Aqua
CD45-
CD45+
HLA-DR – BV421
CD45 – BV605
Big cells
Lymphocytes
SSC-A
FSC-A
CD11b+CD14-
6.76
CD11b+CD14+
82.4
CD11b-CD14-
9.16
CD11b – FITC
CD14 – APC-H7
R10
CXCL10
2.52
IL-6
1.58
LPS + IFNγ
CXCL10
14.2
IL-6
11.6
R10
CXCL10
3.93
IL-6
5.77
LPS + IFNγ
CXCL10
33.1
IL-6
25.7
CXCL10 – PE
IL-6 – Pe-Cy7

#### Table S1. Antiviral drug candidates for entry inhibition of SARS-CoV-2.

| Supplier | Catalog # | Candidate (generic name) | Mechanism of action | Current indications and remarkable properties |
|---|---|---|---|---|
| | | | ***TMPRSS2 inhibitors*** | |
| Selleckchem | S2874 | **Camostat mesylate** | ▪ Inhibits serine proteases [1].<br>▪ Prevents TMPRSS2 to activate SARS-CoV-2 spike protein for viral entry [2]. | ▪ Antiproteinuric drug [1].<br>▪ Approved for pancreatitis in Japan [2]. |
| | | | ***Corticosteroids*** | |
| | | | ➢ Block the initial inflammatory response by inhibiting the production of nitric oxide and eicosanoids that promote vascular permeability and vasodilation. As a result, leukocyte migration to inflamed tissues is decreased [3].<br>➢ Modulate events in chronic inflammation by altering/suppressing T cell activation [3]. | ➢ Immune-mediated diseases, e.g., rheumatoid arthritis, allergies, ulcerative colitis, lupus erythematosus and skin disorders such as psoriasis and dermatitis [3]. |
| Selleckchem | S4430 | **Cortisol** | ▪ Major glucocorticoid in humans [3]. | ▪ It has disadvantageous salt retaining properties which distinguishes it from other glucocorticoids [4]. |
| Sigma-Aldrich | D1756 | **Dexamethasone** | ▪ Possesses 20 to 30 times the binding affinity for glucocorticoid receptors of endogenous cortisol [5]. | ▪ Treatment for post-operative and chemotherapy-induced nausea and vomiting [5]. |
| Selleckchem | S2121 | **Ciclesonide** | ▪ Converted in the lower respiratory tract to an active metabolite with 100-fold greater relative glucocorticoid receptor binding affinity than ciclesonide itself [6 7]. | ▪ Asthma prevention [7]. |
| Selleckchem | S1696 | **Prednisone** | ▪ Metabolized in the liver to its active metabolite [8]. | ▪ First choice in miastenia gravis [8].<br>▪ Maintenance therapy for kidney transplant patients [8]. |
| | | | ***Nonsteroidal anti-inflammatory drugs (NSAIDs)*** | |
| | | | ➢ Inhibit ciclooxygenases (COXs), causing a reduction in the production of prostaglandins at the site of tissue injury and attenuation of the inflammatory cascade [9]. | ➢ Provide analgesia for mild to moderate pain resulting from surgery, injury, and disease [9].<br>➢ Anti-inflammatory properties [9]. |
| Selleckchem | S1261 | **Ibuprofen** | ▪ Inhibits COX [9].<br>▪ Exerts a direct spinal action by blocking the hyperalgesic response induced by activation of spinal glutamate and substance P receptors [9]. | ▪ Widely used as an analgesic, anti-inflammatory, and antipyretic [10]. |
| Selleckchem | S1622 | **Licofelone** | ▪ Inhibits COX and lipoxigenase (LOX) [11]. | ▪ Phase III trials have been successfully completed as a treatment for osteoarthritis[11]. |
| Selleckchem | S1638 | **Sulindac** | ▪ Converted *in vivo* to an active sulfide compound by liver enzymes [12].<br>▪ Inhibits COX [12].<br>▪ Modulates Wnt/β-catenin as well as NF-κB signalling that can have repercussion on the invasive behaviour of cancer cells [13]. | ▪ Relief of signs and symptoms of several arthritic conditions [12].<br>▪ Induction of cholestatic liver injury due to its competitive inhibition of canalicular bile acid transport [14]. |
| Selleckchem | S2386 | **Celecoxib** | ▪ Selectively inhibits COX-2 [15]. | ▪ Pain relief in osteoarthritis [15].<br>▪ Reduction of precancerous polyps in the colon [16].<br>▪ Potentiation of the anticoagulant effects of warfarin. Serious bleeding complications have been reported [17]. |
| | | | ***Other immunomodulators*** | |
| Selleckchem | S4238 | **Cepharanthine** | ▪ Alkaloid isolated from *Stephania cepharantha* [18].<br>▪ Modulates efflux pumps and membrane rigidification [19].<br>▪ Reduces inflammation by AMPK activation and NF-κB inhibition [19].<br>▪ Inhibits plasma membrane lipid peroxidation and platelet aggregation [20].<br>▪ Suppresses cytokine production [20]. | ▪ Used in Japan since the 1950s to treat leukopenia, snake bites, xerostomia and alopecia areata [19].<br>▪ Immunoregulatory, anti-oxidative, -inflammatory, -cancer, -viral and -parasitic properties [19]. |
| Selleckchem | S4008 | **Pemirolast potassium** | ▪ Stabilizes mast cells [21]. | ▪ Prevention and relief of ocular manifestations of allergic conjunctivitis [21]. |

| | | | | |
|---|---|---|---|---|
| | | | ***For enzymatic deficiencies*** | |
| Selleckchem | S4680 | **Protilerin** | ▪ Stimulates the release of thyroid-stimulating hormone from the anterior pituitary gland [22]. | ▪ Stimulation test to diagnose hyperthyroidism [23]. |
| Sigma-Aldrich | T4425 | **Tetrahydrobiopterin dihydrochloride** | ▪ Essential cofactor required for the synthesis of several neurotransmitters [24]. | ▪ Tetrahydrobiopterin deficiency (phenylketonuria) [25]. |
| | | | ***Blood regulation*** | |
| Selleckchem | S5084 | **Carbazochrome sodium sulfonate** | ▪ Reduces capillary permeability [26].<br>▪ Hemostatic agent that promotes clotting [27]. | ▪ Pain relief in refractory chronic prostatitis [26].<br>▪ In combination with tranexamic acid for the reduction of perioperative blood loss and inflammatory response [28].<br>▪ In combination with vitamin C, vitamin E and lysozyme for the reduction of gingival inflammation in chronic periodontitis [29].<br>▪ In combination with procaine results in better efficacy and less adverse effects in treating moderate to massive hemoptysis than vasopressin [30]. |
| MedChemExpress | HY-B0799 | **Ergoloid mesylates** | ▪ Binds with high affinity to the γ-aminobutyric acid $(GABA)_A$ receptor $Cl^-$ channel, producing an allosteric interaction with the benzodiazepine site [31].<br>▪ Dihydrogenation eliminates vasoconstrictor effects of ergotoxine and enhances its α-adrenoreceptor and 5-hydroxytryptamine (serotonin) receptor antagonist properties [32].<br>▪ Inhibits brain-specific phosphodiesterases [32]. | ▪ Supposed therapeutic effects in depression, confusion, lack of self-care in the elderly, and erectile dysfunction [32]. |
| Sigma-Aldrich | SML2313 | **Higenamine hydrochloride** | ▪ Alkaloid found in *Aconitum* plant [33].<br>▪ Reduces IL-1β-induced inflammation in human nucleus pulposus cells via inhibiting NF-κB signaling pathway [34].<br>▪ Protects neuronal cells against oxygen-glucose deprivation/reperfusion (OGD/R)-induced injury by regulating the Akt and Nrf2/HO-1 signaling pathways [35].<br>▪ Possesses positive ionotropic and chronotropic, activating slow channel, vascular and tracheal relaxation effects [33]. | ▪ Collapse, syncope, painful joints, edema, and bronchial asthma treatment in Asian traditional medicine [33].<br>▪ Antispasmodic for Raynaud's phenomenon and cold-induced vasoconstriction [36].<br>▪ Potential therapeutic effects for diseases like intravertebral disc degeneration [34], heart failure, disseminated intravascular coagulation, ischemia/reperfusion injuries and erectile dysfunction [33].<br>▪ Immunomodulatory, anti-inflammatory, -thrombotic, -apoptotic and -oxidative properties [33]. |
| | | | ***Antidiabetics*** | |
| Vall d'Hebron Pharmacy Department | | **Metformin** | ▪ Reduces gluconeogenesis and hepatic glucose production and improves insulin sensitivity by increasing peripheral glucose uptake [37].<br>▪ Increases anaerobic glucose metabolism in enterocytes [38].<br>▪ Accumulates in the mitochondria to inhibit mitochondrial complex I, leading to increased cytoplasmic ADP:ATP and AMP:ATP ratios. These changes activate AMPK to regulate glucose metabolism [38]. | ▪ First choice drug for Type 2 diabetes [39]. |
| Selleckchem | S2542 | **Phenformin** | ▪ Acts on the cell membrane to decrease oxidative phosphorylation [40].<br>▪ Produces tissue anoxia [40].<br>▪ Increases peripheral glucose uptake (Pasteur Effect) [40].<br>▪ Leads to lactic acidosis by inhibition of lactic acid metabolism [40]. | ▪ Predecessor of metformin with an unacceptably high incidence of lactic acidosis, often fatal [41]. |
| Vall d'Hebron Pharmacy Department | | **Vildagliptin** | ▪ Selectively inhibits dipeptidyl peptidase-4 (DPP-4), an enzyme that degrades and inactivates glucagon-like peptide-1 (GLP-1) and glucose-dependent insulinotropic polypeptide (GIP), which promote insulin secretion and regulate blood glucose levels [42]. | ▪ Type 2 diabetes [43]. |
| Vall d'Hebron Pharmacy Department | | **Sitagliptin** | ▪ Selectively inhibits DPP-4 [43].<br>▪ Elevates GLP-1 levels to increase insulin release after meals and improve glucose tolerance [43]. | ▪ Type 2 diabetes [43]. |

| | | | | |
|---|---|---|---|---|
| | | | ***Natural compounds - Flavonoids*** | |
| | | | ➢Possess several anticancer effects: they modulate reactive oxygen species (ROS)-scavenging enzyme activities, participate in arresting the cell cycle, induce apoptosis, autophagy, and suppress cancer cell proliferation and invasiveness [44]. | ➢Multiple potential applications. |
| Selleckchem | S2320 | **Luteolin** | ▪ Found in a number of dietary sources [45]. | ▪ Plants rich in luteolin have been used in Chinese traditional medicine for hypertension, inflammatory disorders, and cancer treatment [46]. |
| Sigma-Aldrich | 94258 | **Eriodictyol** | ▪ Present in citrus fruits and Chinese herbs used in the food industry [47].<br>▪ Reduces inflammation by NF-κB blockade [48].<br>▪ As opposite to other flavonoids, eriodictyol lacks the C2–C3 double bond responsible to give more inhibitory activity of basal vascular NO release and vascular superoxide formation [49] | ▪ Anti-inflammatory, -allergenic, -microbial, -cancer, and -oxidant properties [48]. |
| Selleckchem | S2391 | **Quercetin** | ▪ Obtained from diverse fruits and vegetables [50].<br>▪ Stabilizes basophils and mast cells [50].<br>▪ Reduces obesity by increase in AMPK expression [51].<br>▪ Inhibits histone deacetylase 1 and DNA methyltransferase 1 [51]. | ▪ Antioxidative, -inflammatory, -platelet, -apoptotic, -invasive and -angiogenic properties [51].<br>▪ Nephro-, gastro-, angio-, cardio- and chondroprotective properties [52]. |
| Selleckchem | S2007 | **Myricetin** | ▪ Found in tea, berries, fruits, vegetables, and the plant *Diospyros lotus* [53].<br>▪ Inhibits thrombin with an $IC_{50}$ value of 56 μM [54].<br>▪ Inhibits the production of proinflammatory mediators through the suppression of NF-κB and STAT1 activation and induction of Nrf2-mediated HO-1 expression in LPS-stimulated RAW264.7 macrophages [55]. | ▪ *Diospyros lotus* is traditionally used in diabetes, diarrhea, tumor, and hypertension treatment [55]. |
| | | | ***Natural compounds - Others*** | |
| Selleckchem | S2326 | **Glycyrrhizin** | ▪ Triterpenoid saponin found as the major active constituent in licorice root [56].<br>▪ Shows anti-inflammatory and antioxidant activities [57].<br>▪ Stimulates endogenous production of interferons [57].<br>▪ Induces melanogenesis through cAMP signaling [58]. | ▪ Bronchitis, gastritis, and jaundice alleviation in traditional medicine [56]. |
| Selleckchem | S4646 | **Indirubin** | ▪ Found in a variety of plants, marine mollusks, bacteria, and human urine, and produced in mammals' intestine [59].<br>▪ Inhibits several cyclin-dependent kinases and glycogen synthase kinase 3 [59].<br>▪ May suppress lipopolysaccharides-induced inflammation via Toll-like receptor 4 abrogation mediated by the NF-κB and MAPK signaling pathways [60]. | ▪ Chronic myelogenous leukemia treatment in Chinese traditional medicine [59].<br>▪ Indirubin in Lindioil ointment for psoriasis topical treatment [61]. |
| MedChemExpress | HY-N0453 | **Hypericin** | ▪ Produces primary photosensitization [62].<br>▪ Shows selective activity against viruses, both *in vitro* and *in vivo*, including Herpes simplex (HSV)-1 and -2 [63]. | ▪ Non-melanoma skin cancer topical treatment [62]. |
| | | | ***Antimicrobials*** | |
| Sigma-Aldrich | 90527 | **Hydroxychloroquine** | ▪ Inhibits pH-dependent viral fusion/replication [1].<br>▪ Prevents viral envelope glycoprotein as well as host receptor protein glycosylation [1].<br>▪ Prevents virion assembly in endoplasmic reticulum-Golgi intermediate compartment–like structure [1].<br>▪ Inhibits TLR-7/9–dependent inflammatory responses [1]. | ▪ Malaria [64].<br>▪ Rheumatoid arthritis [64].<br>▪ Systemic lupus erythematosus [64]. |
| Sigma-Aldrich | M2140 | **Monocaprin** | ▪ Safe functional emulsifier in food industry [65].<br>▪ May be a potential preservative independent of pH, acting by disruption of the cell wall and plasma membrane of fungi [65]. | ▪ Hydrogel formulations have *in vitro* microbicidal activity against HIV and HSV, *Chlamydia trachomatis* and *Neisseria gonorrhoeae* [66].<br>▪ In combination with doxycycline as a potential treatment of herpes labialis [67]. |
| Sigma-Aldrich | I8898 | **Ivermectin** | ▪ GABA receptor agonist [68].<br>▪ Exerts toxicity in parasites by blocking the post-synaptic transmission of nerve impulses [68]. | ▪ Broad-spectrum anti-parasite medication [69].<br>▪ First choice treatment for onchocerciasis [69]. |

| | | | | |
|---|---|---|---|---|
| Vall d'Hebron Pharmacy Department | | **Nitrofurantoin** | ▪ Bacterial intracellular nitroreductases produce the active form of the drug [70].<br>▪ Intermediate metabolites bind to bacterial ribosomes and inhibit bacterial enzymes involved in the synthesis of DNA, RNA, and bacterial wall protein synthesis [70]. | ▪ Uncomplicated urinary tract infections resistant to other antibiotics [71]. |
| Fisher Scientific | 10387340 | **Lauric acid** | ▪ Found in vegetal oils [72].<br>▪ Disrupts the cell membrane of gram-positive bacteria by physicochemical processes [72].<br>▪ Interferes with bacterial cell signal transduction and gene transcription processes [72].<br>▪ Activates TLR4 signaling [73]. | ▪ Main antiviral and antibacterial substance found in human breast milk [74]. |
| Selleckchem | S3132 | **Sulfamethoxazole** | ▪ A sulfonamide derivative. Sulfonamides have a bacteriostatic effect by inhibiting bacterial folic acid synthesis [75]. | ▪ In combination with trimethoprim for urinary tract infections, otitis media, chronic bronchitis, *Shigella* and enterotoxigenic *Escherichia coli* infections [76].<br>▪ Prophylaxis of *Pneumocystis jirovecii* pneumonia in patients with HIV/AIDS [77].<br>▪ High rates of adverse drug reactions in subjects infected by HIV [78]. |
| Selleckchem | S2302 | **Sulfamerazine** | ▪ A sulfonamide derivative [75]. | ▪ Bronchitis, prostatitis, and urinary tract infections [79].<br>▪ Combined with a folate antagonist, sulfonamides are indicated among others in toxoplasmosis and malaria [80]. |
| Vall d'Hebron Pharmacy Department | | **Tazobactam** | ▪ Class A β-lactamase inhibitor [81]. | ▪ Extension of antibiotic's spectrum of activity [81].<br>▪ Increase in β-lactamic antibiotic stability against bacterial β-lactamases [81]. |
| Selleckchem | S1784 | **Vidarabine** | ▪ Purine nucleoside analogue [80].<br>▪ Competitively inhibits DNA-dependent DNA polymerases of some DNA viruses approximately 40 times more than those of host cells [80]. | ▪ The degree of maximal resistance to vidarabine is 4-fold, much lower than the 100-fold resistance to acyclovir with similar DNA-polymerase resistant mutations [80].<br>▪ No longer used due to toxicity issues as well as the discovery of more potent and safer compounds such as acyclovir, which today is widely prescribed for HSV [82]. |
| Sigma-Aldrich | M1765 | **Monolaurin** | ▪ Solubilizes the lipids and phospholipids in the envelope of pathogenic viruses and bacteria causing the disintegration of their envelopes [74]. | ▪ Inactivation to some extent of HIV, measles, HSV-1, vesicular stomatitis, Visna virus, and cytomegalovirus [74]. |
| Sigma-Aldrich | PHR1949 | **Sodium lauryl sulfate** | ▪ Ionic detergent that rapidly disrupts biological membranes [83]. | ▪ Inhibition of the formation of several bacterial biofilms [84]. |
| Selleckchem | S1915 | **Valaciclovir** | ▪ Purine nucleoside analogue [80].<br>▪ Rapidly and extensively converted to aciclovir by first-pass metabolism [85]. | ▪ Highly active against Herpes simplex and Herpes zoster [85]. |

#### REFERENCES


1. Hosoki K, Chakraborty A, Sur S. Molecular mechanisms and epidemiology of COVID-19 from an allergist's perspective. *Journal of Allergy and Clinical Immunology.* 2020;146(2):285-299.
2. Hoffmann M, Hofmann-Winkler H, Smith JC, et al. Camostat mesylate inhibits SARS-CoV-2 activation by TMPRSS2-related proteases and its metabolite GBPA exerts antiviral activity. *EBioMedicine.* Mar 2021;65:103255.
3. Cruz-Topete D, Cidlowski JA. Glucocorticoids: Molecular Mechanisms of Action. In: Riccardi C, Levi-Schaffer F, Tiligada E, eds. *Immunopharmacology and Inflammation*. Cham: Springer International Publishing; 2018:249-266.
4. Hindmarsh PC, Geertsma K. Chapter 20 - Hydrocortisone. In: Hindmarsh PC, Geertsma K, eds: Academic Press; 2017:231-249.
5. Zabirowicz ES, Gan TJ. 34 - Pharmacology of Postoperative Nausea and Vomiting. In: Hemmings HC, Egan TD, eds. Second Edition ed. Philadelphia: Elsevier; 2019:671-692.
6. Ciclesonide. In: Aronson JK, ed. *Meyler's Side Effects of Drugs*. Sixteenth Edition ed. Oxford: Elsevier; 2016:292-294.
7. Kercsmar CM, McDowell KM. 45 - Wheezing in Older Children: Asthma. In: Wilmott RW, Deterding R, Li A, et al., eds. Ninth Edition ed. Philadelphia: Elsevier; 2019:686-721.e684.
8. Silvestri NJ, Barohn RJ, Wolfe GI. 145 - Acquired Disorders of the Neuromuscular Junction. In: Swaiman KF, Ashwal S, Ferriero DM, et al., eds. Sixth Edition ed: Elsevier; 2017:1098-1105.
9. Walker BJ, Polaner DM, Berde CB. 44 - Acute Pain. In: Coté CJ, Lerman J, Anderson BJ, eds. Sixth Edition ed. Philadelphia: Elsevier; 2019:1023-1062.e1015.
10. DrugBank. Ibuprofen. https://go.drugbank.com/drugs/DB01050.
11. Information NCfB. PubChem Compound Summary for CID 133021, Licofelone. 2021.
12. DrugBank. Sulindac. https://go.drugbank.com/drugs/DB00605.
13. Sherbet GV. Chapter 25 - S100A4 Has Potential Benefits as a Therapeutic Target. In: Sherbet GV, ed: Academic Press; 2017:211-221.
14. Manautou JE, Campion SN, Aleksunes LM. 9.08 - Regulation of Hepatobiliary Transporters during Liver Injury. In: McQueen CA, ed. Second Edition ed. Oxford: Elsevier; 2010:175-220.
15. Roy J. 11 - Life-style drugs, statins, and COX-2 drugs. In: Roy J, ed: Woodhead Publishing; 2011:297-325.
16. DrugBank. Celecoxib. https://go.drugbank.com/drugs/DB00482.
17. Celecoxib. In: Aronson JK, ed. *Meyler's Side Effects of Drugs*. Sixteenth Edition ed. Oxford: Elsevier; 2016:191-195.
18. Nair A, Chattopadhyay D, Saha B. Chapter 17 - Plant-Derived Immunomodulators. In: Ahmad Khan MS, Ahmad I, Chattopadhyay D, eds: Academic Press; 2019:435-499.
19. Bailly C. Cepharanthine: An update of its mode of action, pharmacological properties and medical applications. *Phytomedicine.* 2019;62:152956-152956.
20. Rogosnitzky M, Danks R. Therapeutic potential of the biscoclaurine alkaloid, cepharanthine, for a range of clinical conditions. *Pharmacological Reports.* 2011;63(2):337-347.
21. Bielory L, Bielory BP, Wagner RS. 54 - Allergic and Immunologic Eye Disease. In: Leung DYM, Szefler SJ, Bonilla FA, Akdis CA, Sampson HA, eds. Third Edition ed. London: Elsevier; 2016:482-497.e483.
22. Pirahanchi Y, Toro F, Jialal I. Physiology, Thyroid Stimulating Hormone. 2020; https://www.ncbi.nlm.nih.gov/books/NBK499850/.

23. Manolov D, Zakharieva B, Lazarov G, Manov A, Vŭrbanov V. The TRH test using the Bulgarian preparation Protilerin in the diagnosis of hyperthyroidism. *Vutreshni bolesti.* 1989;28(5):30-33.
24. Bhagavan NV, Ha C-E. Chapter 15 - Protein and Amino Acid Metabolism. In: Bhagavan NV, Ha C-E, eds. *Essentials of Medical Biochemistry*. Second Edition ed. San Diego: Academic Press; 2015:227-268.
25. Muntau AC, Adams DJ, Bélanger-Quintana A, et al. International best practice for the evaluation of responsiveness to sapropterin dihydrochloride in patients with phenylketonuria. *Molecular Genetics and Metabolism.* 2019;127(1):1-11.
26. Oh-oka H, Yamada T, Noto H, et al. Effect of carbazochrome sodium sulfonate on refractory chronic prostatitis. *International Journal of Urology.* 2014;21(11):1162-1166.
27. DrugBank. Carbazochrome. https://go.drugbank.com/drugs/DB09012.
28. Luo Y, Zhao X, Releken Y, Yang Z, Pei F, Kang P. Hemostatic and Anti-Inflammatory Effects of Carbazochrome Sodium Sulfonate in Patients Undergoing Total Knee Arthroplasty: A Randomized Controlled Trial. *The Journal of Arthroplasty.* 2020;35(1):61-68.
29. Hong J-Y, Lee J-S, Choi S-H, et al. A randomized, double-blind, placebo-controlled multicenter study for evaluating the effects of fixed-dose combinations of vitamin C, vitamin E, lysozyme, and carbazochrome on gingival inflammation in chronic periodontitis patients. *BMC Oral Health.* 2019;19(40).
30. Ren CS, Du YP, Yu XY. The effect of Procaine combined with Carbazochrome on moderate to massive hemoptysis due to tuberculosis. *Med. J. West China.* 2011;23:1045-1046.
31. MedChemExpress. Dihydroergotoxine mesylate. https://www.medchemexpress.com/Dihydroergotoxine_mesylate.html.
32. Ergot derivatives. In: Aronson JK, ed. *Meyler's Side Effects of Drugs*. Sixteenth Edition ed. Oxford: Elsevier; 2016:86-92.
33. Zhang N, Lian Z, Peng X, Li Z, Zhu H. Applications of Higenamine in pharmacology and medicine. *Journal of Ethnopharmacology.* 2017;196:242-252.
34. Bai X, Ding W, Yang S, Guo X. Higenamine inhibits IL-1β-induced inflammation in human nucleus pulposus cells. *Bioscience Reports.* 2019;39(6).
35. Zhang Y, Zhang J, Wu C, et al. Higenamine protects neuronal cells from oxygen-glucose deprivation/reoxygenation-induced injury. *Journal of Cellular Biochemistry.* 2019;120(3):3757-3764.
36. Guan J, Lin H, Xie M, et al. Higenamine exerts an antispasmodic effect on cold-induced vasoconstriction by regulating the PI3K/Akt, ROS/α2C-AR and PTK9 pathways independently of the AMPK/eNOS/NO axis. *Experimental and Therapeutic Medicine.* 2019;18(2):1299-1308.
37. Day C. Chapter 26 - New Therapies in Obesity. In: Weaver JU, ed: Elsevier; 2018:271-279.
38. DrugBank. Metformin. https://go.drugbank.com/drugs/DB00331.
39. Brindle PK. Chapter 253 - Transcriptional Regulation via the cAMP Responsive Activator CREB. In: Bradshaw RA, Dennis EA, eds. Second Edition ed. San Diego: Academic Press; 2010:2077-2081.
40. Vaman Rao C. Biguanides. In: Wexler P, ed. Second Edition ed. New York: Elsevier; 2005:271-272.
41. DrugBank. Phenformin. https://go.drugbank.com/drugs/DB00914.
42. Sonia TA, Sharma CP. 1 - Diabetes mellitus – an overview. In: Sonia TA, Sharma CP, eds: Woodhead Publishing; 2014:1-57.
43. Bennett RG. Sitagliptin. *Reference Module in Biomedical Sciences*: Elsevier; 2018.
44. Kopustinskiene DM, Jakstas V, Savickas A, Bernatoniene J. Flavonoids as anticancer agents. *Nutrients.* 2020;12(2):457-457.
45. Omar SH. Chapter 4 - Biophenols: Impacts and Prospects in Anti-Alzheimer Drug Discovery. In: Brahmachari G, ed. *Discovery and Development of Neuroprotective Agents from Natural Products*: Elsevier; 2018:103-148.
46. Avendaño C, Menéndez JC. Cancer Chemoprevention. *Medicinal Chemistry of Anticancer Drugs*. Second Edition ed: Elsevier; 2015:701-723.

47. Rajan VK, Muraleedharan K, Hussan KPS. Structural Evaluation and Toxicological Study of a Bitter Masking Bioactive Flavanone, 'Eriodictyol'. *Polyphenols: Prevention and Treatment of Human Disease*. Second Edition ed: Elsevier; 2018:45-60.
48. Shukla R, Pandey V, Vadnere GP, Lodhi S. Chapter 18 - Role of Flavonoids in Management of Inflammatory Disorders. In: Watson RR, Preedy VR, eds. *Bioactive Food as Dietary Interventions for Arthritis and Related Inflammatory Diseases*. Second Edition ed: Academic Press; 2019:293-322.
49. Ribeiro D, Fernandes E, Freitas M. Flavonoids as Modulators of Neutrophils' Oxidative Burst: Structure-Activity Relationship. Elsevier; 2018:261-276.
50. Horwitz RJ. The Allergic Patient. Elsevier; 2018:300-309.e302.
51. Woon ECY, Toh JDW. Chapter 6 - Antiobesity Effects of Natural Products from an Epigenetic Perspective. *Studies in Natural Products Chemistry.* Vol 41, 2014:161-193.
52. Shebeko SK, Zupanets IA, Popov OS, Tarasenko OO, Shalamay AS. Effects of Quercetin and Its Combinations on Health. Elsevier; 2018:373-394.
53. Jadeja RN, Devkar RV. Polyphenols and Flavonoids in Controlling Non-Alcoholic Steatohepatitis. Elsevier; 2014:615-623.
54. Wang X, Yang Z, Su F, et al. Study on structure activity relationship of natural Flavonoids against thrombin by molecular docking virtual screening combined with activity evaluation *in vitro*. *Molecules.* 2020;25(2):422-422.
55. Cho BO, Yin HH, Park SH, Byun EB, Ha HY, Jang SI. Anti-inflammatory activity of myricetin from *Diospyros lotus* through suppression of NF-κB and STAT1 activation and Nrf2-mediated HO-1 induction in lipopolysaccharide-stimulated RAW264.7 macrophages. *Bioscience, Biotechnology, and Biochemistry.* 2016;80(8):1520-1530.
56. Girish C, Pradhan SC. Herbal Drugs on the Liver. Elsevier; 2017:605-620.
57. Ramos-Tovar E, Muriel P. Phytotherapy for the Liver. In: Ronald Ross Watson VRP, ed. *Dietary Interventions in Liver Disease*: Academic Press; 2019:101-121.
58. Blom van Staden A, Lall N. Chapter 5 - Medicinal Plants as Alternative Treatments for Progressive Macular Hypomelanosis. *Medicinal Plants for Holistic Health and Well-Being*: Elsevier; 2018:145-182.
59. Cragg GM, Newman DJ. Natural Product Sources of Drugs: Plants, Microbes, Marine Organisms, and Animals. Elsevier; 2007:355-403.
60. Lai J-l, Liu Y-h, Liu C, et al. Indirubin inhibits LPS-induced inflammation via TLR4 abrogation mediated by the NF-kB and MAPK signaling pathways. *Inflammation.* 2017;40(1):1-12.
61. Lin YK, See LC, Huang YH, Chi CC, Hui RCY. Comparison of indirubin concentrations in indigo naturalis ointment for psoriasis treatment: a randomized, double-blind, dosage-controlled trial. *British Journal of Dermatology.* 2018;178(1):124-131.
62. Penjweini R, Deville S, Ethirajan A, Ameloot M. Chapter 22 - Investigating the Intracellular Dynamics of Hypericin-Loaded Nanoparticles and Polyvinylpyrrolidone-Hypericin by Image Correlation Spectroscopy. In: Hamblin MR, Avci P, Prow TW, eds. Boston: Academic Press; 2016:275-286.
63. Romm A, Clare B, Alschuler L, Hobbs C, Upton R. Chapter 8 - Vaginal Infections and Sexually Transmitted Diseases. In: Romm A, Hardy ML, Mills S, eds. Saint Louis: Churchill Livingstone; 2010:256-289.
64. Howard B. Hydroxychloroquine. In: S.J. Enna DBB, ed. *xPharm: The Comprehensive Pharmacology Reference*: Elsevier; 2007:1-4.
65. Ma M, Wen X, Xie Y, et al. Antifungal activity and mechanism of monocaprin against food spoilage fungi. *Food Control.* 2018;84:561-568.
66. Neyts J, Kristmundsdottir T, De Clercq E, Thormar H. Hydrogels containing monocaprin prevent intravaginal and intracutaneous infections with HSV-2 in mice: Impact on the search for vaginal microbicides. *Journal of Medical Virology.* 2000;61(1):107-110.
67. Skulason S, Holbrook WP, Thormar H, Gunnarsson GB, Kristmundsdottir T. A study of the clinical activity of a gel combining monocaprin and doxycycline: a novel treatment for herpes labialis. *Journal of Oral Pathology & Medicine.* 2012;41(1):61-67.
68. Gupta RC, Milatovic D. Chapter 23 - Insecticides. In: Gupta RC, ed. *Biomarkers in Toxicology*. Boston: Academic Press; 2014:389-407.

69. Bogitsh BJ, Carter CE, Oeltmann TN. Chapter 17 - Blood and Tissue Nematodes. In: Burton J. Bogitsh CEC, Thomas N. Oeltmann, ed. *Human Parasitology*. Fifth Edition ed: Academic Press; 2019:313-329.
70. Squadrito FJ, del Portal D. Nitrofurantoin. *StatPearls*. Treasure Island (FL)2021.
71. Howard B, Furman B. Nitrofurantoin. *Reference Module in Biomedical Sciences*: Elsevier; 2018.
72. Dayrit FM. The properties of lauric acid and their significance in coconut oil. *Journal of the American Oil Chemists' Society.* 2015;92(1):1-15.
73. Wang L, Luo L, Zhao W, et al. Lauric Acid Accelerates Glycolytic Muscle Fiber Formation through TLR4 Signaling. *Journal of agricultural and food chemistry.* 2018;66(25):6308-6316.
74. Lieberman S, Enig MG, Preuss HG. A review of monolaurin and lauric acid: natural virucidal and bactericidal agents. *Alternative and Complementary Therapies.* 2006;12(6):310-314.
75. Singh AK, Sharma AK, Khan I, Gothwal A, Gupta L, Gupta U. Chapter 9 - Oral drug delivery potential of dendrimers. In: Andronescu E, Grumezescu AM, eds. *Nanostructures for Oral Medicine*: Elsevier; 2017:231-261.
76. DrugBank. Sulfamethoxazole. https://go.drugbank.com/drugs/DB01015.
77. Walker L, Yip V, Pirmohamed M. Chapter 20 - Adverse Drug Reactions. In: Padmanabhan S, ed. San Diego: Academic Press; 2014:405-435.
78. Antifungal azoles [for systemic use]. In: Aronson JK, ed. *Meyler's Side Effects of Drugs*. Sixteenth Edition ed. Oxford: Elsevier; 2016:584-601.
79. DrugBank. Sulfamerazine. https://go.drugbank.com/drugs/DB01581.
80. Padberg S. 2.6 - Anti-infective Agents. In: Schaefer C, Peters P, Miller RK, eds. Third Edition ed. San Diego: Academic Press; 2015:115-176.
81. Kuriyama T, Karasawa T, Williams DW. Chapter 13 - Antimicrobial Chemotherapy: Significance to Healthcare. In: Percival SL, Williams DW, Randle J, Cooper T, eds. *Biofilms in Infection Prevention and Control*. Boston: Academic Press; 2014:209-244.
82. Seley-Radtke KL, Yates MK. The evolution of nucleoside analogue antivirals: A review for chemists and non-chemists. Part 1: Early structural modifications to the nucleoside scaffold. *Antiviral Research.* 2018;154:66-86.
83. Farrell RE. Chapter 7 - Resilient Ribonucleases. In: Farrell RE, ed. *RNA Methodologies*. Fourth Edition ed. San Diego: Academic Press; 2010:155-172.
84. Shen Y, Li P, Chen X, et al. Activity of Sodium Lauryl Sulfate, Rhamnolipids, and N-Acetylcysteine against biofilms of five common pathogens. *Microbial Drug Resistance.* 2020;26(3):290-299.
85. Valaciclovir. In: Aronson JK, ed. *Meyler's Side Effects of Drugs*. Sixteenth Edition ed. Oxford: Elsevier; 2016:297-300.

**Table S2. $EC_{50}$ and $CC_{50}$ of 39 antiviral drug candidates.**

| Drugs | Vero E6 | HLT | Vero E6 | HLT | |
|---|---|---|---|---|---|
| | $EC_{50}$ (μM) | | $CC_{50}$ (μM) | | |
| Cepharanthine | 0.46 | 6.08 | 22.37 | 16.15 | Concordant |
| Luteolin | ~70.70 | ~82.39 | >100 | >100 | |
| Ergoloid | 4.78 | 9.17 | 18.67 | ~100 | |
| Ciclesonide | 20.52 | 16.41 | >100 | >100 | |
| Licofelone | 87.50 | 32.16 | >100 | ~53.61 | |
| Hydroxychloroquine | 1.58 | 3.22 | >100 | ~100 | |
| Ivermectin | 13.94 | 12.98 | 20.23 | ~100 | |
| Celecoxib | 12.99 | 55.72 | 12.98 | ~100 | |
| Hypericin | 1.24 | 0.31 | 4.80 | 0.14 | |
| Vidarabine | >100 | 51.60 | >100 | >100 | |
| Eriodictyol | No effect | 90.00 | >100 | >100 | Discordant |
| Quercetin | >100 | No effect | >100 | >100 | |
| Camostat | No effect | 3.30 | >100 | >100 | |
| Phenformin | No effect | 38.16 | >100 | >100 | |
| Valaciclovir | No effect | >100 | >100 | >100 | |
| Sulindac | No effect | >100 | >100 | >100 | |
| SLS | ~26.45 | >100 | 87.05 | ~83.2 | No effect |
| Myricetin | No effect | No effect | >100 | >100 | |
| Sitagliptin | No effect | No effect | >100 | >100 | |
| Dexamethasone | No effect | No effect | >100 | >100 | |
| Pemirolast | No effect | No effect | >100 | >100 | |
| Protilerin | No effect | No effect | >100 | >100 | |
| Sulfamerazine | No effect | No effect | >100 | >100 | |
| Carbazochrome | No effect | No effect | >100 | >100 | |
| Higenamine | No effect | No effect | >100 | >100 | |
| Tetrahydrobiopterin | No effect | No effect | >100 | >100 | |
| Metformin | No effect | No effect | >100 | >100 | |
| Vidagliptin | No effect | No effect | >100 | >100 | |
| Monocaprin | No effect | No effect | >100 | >100 | |
| Lauric acid | No effect | No effect | >100 | >100 | |
| Monolaurin | No effect | No effect | >100 | >100 | |
| Tazobactam | No effect | No effect | >100 | >100 | |
| Nitrofurantoin | No effect | No effect | >100 | >100 | |
| Cortisol | No effect | No effect | >100 | >100 | |
| Ibuprofen | No effect | No effect | >100 | >100 | |
| Indirubin | No effect | No effect | >100 | >100 | |
| Glycyrrhizin | No effect | No effect | >100 | >100 | |
| Sulfamethoxazole | No effect | No effect | >100 | >100 | |
| Prednisone | No effect | Some effect | >100 | >100 | |
